# Long-term responses of Icelandic Arctic foxes to changes in marine and terrestrial ecosystems

**DOI:** 10.1101/2023.02.09.527803

**Authors:** Fanny Berthelot, Ester R. Unnsteinsdóttir, Jennifer A. Carbonell Ellgutter, Dorothee Ehrich

## Abstract

The long-term dynamics of predator populations may be driven by fluctuations in resource availability and reflect ecosystem changes such as those induced by climate change. The Icelandic Arctic fox (*Vulpes lagopus*) population has known major fluctuations in size since the 1950s. Using stable isotopes analysis of bone collagen over a long-time series (1979-2018), we aimed at identifying the main resources used by Icelandic Arctic foxes during periods of growth and decline to assess if the variations in their population size are linked to fluctuations in the availability of resources. We hypothesized that (1) the decline in Seabirds abundance was responsible for the decrease in the fox population; and (2) that the growth in the fox population combined to fluctuations in main resources would lead to an increase in intra-specific competition, ultimately leading to variations in their isotopic niches at the population scale. The isotopic signatures of Arctic foxes as well as their temporal trends differed clearly between inland and coast. Inland foxes showed an overall shift towards more terrestrial preys, whereas coastal foxes displayed a relatively stable use of marine resources over the years. Stable isotopes mixing models suggested that marine resources and rock ptarmigans were the most important food source and highlighted a rather stable diet in coastal habitats compared to inland habitats where more fluctuations in dietary composition were observed. Coastal foxes had a broader niche than inland foxes, and there was more variation in niche size in the inland habitat. Our results are in agreement with a general decline in seabird populations driving the decline in Arctic foxes, especially in coastal habitats. For the inland foxes, our results suggest that the lack of marine resources might have led to an increased use of ptarmigans as well as other terrestrial resources such as geese and waders, especially during the most recent period.

## Introduction

Many Arctic ecosystems are characterized by a close link to adjacent marine environments. Around 80% of the terrestrial Arctic lays within 100 km from the coast [1], and marine subsidies often play an important role in terrestrial food webs [2,3]. At present, high latitude ecosystems are subject to rapid changes under the influence of global warming [4,5]. These changes may affect marine and terrestrial systems differently and thus modify the interplay between these neighboring ecosystems [6]. Warmer and longer summers may lead to increased primary production on land, and better conditions for some herbivores [7], whereas expanding boreal species can become serious competitors of arctic endemics [8]. Warming of the sea causes major changes in food webs and leads notably to declines of many seabird populations [9]. Several predator species, such as gulls (*Larus* spp.), skuas (*Stercorarius* spp.) or Arctic foxes (*Vulpes lagopus*) are able to exploit both marine and terrestrial resources, thereby constituting a link between both systems [3,6]. As these opportunistic species are likely to reflect changes through complex ecological responses, their long-term monitoring is a challenging yet key process for a better understanding of the impacts of global warming on interconnected marine and terrestrial ecosystems [10].

The Arctic fox is a terrestrial mammalian predator endemic to the Arctic tundra. It has a circumpolar distribution and is abundant across most of its range, including most arctic islands. Different threats such as increasing competition with the red fox (*Vulpes vulpes*), habitat loss arising from climate change and declines in the abundance of key prey make Arctic foxes increasingly vulnerable in part of their range [11,12]. Therefore, they have been chosen as a climate change flagship species by the International Union of Conservation of Nature [13]. They are also a target of international monitoring as their reliance on tundra ecosystems make them likely to highlight the impacts of climate change through species interactions [10,14].

Arctic foxes have been attributed to two different resource use strategies that involve different reproductive patterns: lemming foxes and coastal foxes [15–17]. The first type behaves as an opportunistic lemming (*Lemmus and Dicrostonyx spp.*) specialist and adapts its breeding effort to the lemming cycle with large litters in peak years [3,17,18]. The second type is more generalist, and lives on Arctic islands deprived of lemmings such as Iceland, or Svalbard. These foxes feed on both marine and terrestrial resources [16,19] and dispose of a more stable food supply, thus producing fewer cubs per year, but breeding more regularly [17]. While the population dynamics of lemming foxes primarily follow the cycles of their prey, coastal foxes are driven by trends in both terrestrial and marine resources [6,17,20].

Iceland makes up for a particularly interesting system when it comes to understanding Arctic fox population dynamics for several reasons. First, this specific coastal population is neither threatened by interspecific competition with the red fox, as Arctic foxes are the only canid species living in Iceland, nor by the collapse of rodent cycles because lemmings are absent from this island [19]. Second, the species being considered as a pest, hunting is known to be the main cause of mortality [21]. The hunting pressure is thought to be stable since 1950, and is regulated by Icelandic laws, which makes it an unlikely driver of the population dynamics, but the hunting statistics provide a long-term estimate of the population trends [20,22]. The island is also free of infectious diseases that could potentially be fatal to foxes, such as rabies or distemper [23,24].

Despite the stable hunting effort and the apparent absence of common ecological pressures, striking long-term fluctuations in Arctic fox numbers have been documented and attributed to variations in carrying capacity, likely driven by the distribution and fluctuations in abundance of prey [20,22]. In addition, [25] found evidence for indirect climatic impacts through food availability. The hunting statistics indicated a decrease of the population from 1950 to 1970, which has been partly explained by a reduction in the rock ptarmigan (*Lagopus muta*) population [20,26,27]. This period of decline has been followed by a steady six-fold increase until 2008 which has been explained by a global rise in food abundance. [25] suggested that climatic variables such as the Sub-Polar Gyre, the North Atlantic Oscillation and summer temperature acted indirectly to increase the abundance of the main preys. Based on prey remains at dens, [20] documented an increased predation on waders and geese during this growth period, as well as an important use of fulmars. Using stable isotopes analysis, [28] highlighted the importance of marine resources and suggested that they might have supported the increase in the fox population. Their results did however not support an increased consumption of geese as suggested by [20].

Recent population estimates have shown important fluctuations in foxes’ numbers during the last decade, starting with a drastic drop reducing the population to half its size within 5 years (Fig 1). Since 2011, however, the population seems to recover. Stable isotopes of carbon and nitrogen reflect the main resources used by a consumer over a certain period of time and can thus provide a good insight into potential drivers of population changes [29]. Building on the study of [28], we used isotopic signatures of bone collagen over 40 years to identify the main resources used by Icelandic Arctic foxes during periods of population growth and decline. We investigate whether the recent fluctuations in the size of the population were associated with changes in main dietary components and assess whether they can be attributed to fluctuations in the availability of prey. As [28] showed that the long growth in population was characterized by a rather constant diet and may thus have been driven by increasing seabird populations, that constitute a key resource, we hypothesized that the recent decline in several seabird species [9] might have negatively affected the foxes during the last decade, especially in coastal areas. In the inland, population fluctuations may be related to a switch in diet from rock ptarmigan to increased predation on geese [20]. The important growth of the fox population together with fluctuations in main resources may have also led to an increase in intra-specific competition. [22] showed that density dependence is one of the main driver of Icelandic Arctic fox population, pointing out that foxes adapt their territory sizes in response to variations in carrying capacity. Therefore, we hypothesized an increase in inter-individual variability in the diet with increasing population size, potentially leading to variation in the niche breadth at the population level. A better understanding of what caused the important variations this population experienced in recent years will be important for further management and hunting recommendations, and shed new light on how the interplay between changes in the marine and terrestrial ecosystems affect a flexible arctic predator.

**Fig 1.**
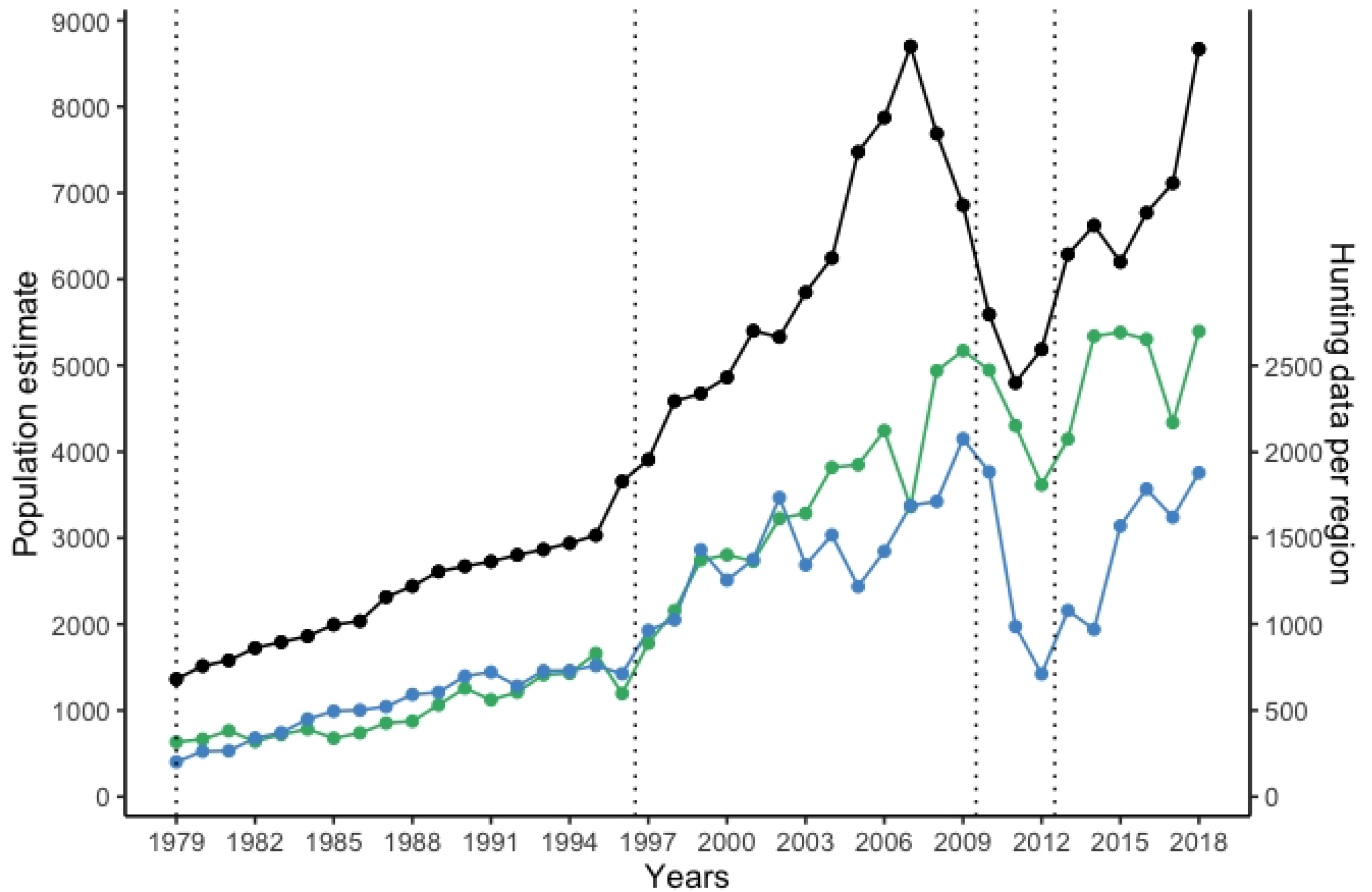
Icelandic Arctic fox population estimates and hunting data per region from 1979 to 2018. The Icelandic Arctic fox population estimates (in black) is based on age cohort analysis and hunting statistics [30]. Foxes culled further than 50km from the coast were referred as inland, while foxes culled within 5km from the coast were considered coastal. The dotted lines delimitate the different period of growth and decline we are using in the study: 1979-1996; 1997-2009; 2010-2012; 2013-2018.

## Material and methods

### Study area and species

Iceland is located in the Atlantic Ocean, close to the Arctic Circle. Its climate is influenced by the Gulf Stream and temperature is considerably higher than might be expected at this latitude. Monthly mean temperatures vary from −3 to 3°C in January and reach 8 to 15°C in July. Precipitation is high, ranging from 400 to 4000 mm annually (climateknowledgeportal.worldbank.org). The temperatures usually prevent the shorelines from freezing throughout the year. Mean annual temperature has on average been above 2°C during the last decades, whereas it was below that during the last century. Iceland is also free from pack-ice and thereby remains isolated [31].

Icelandic Arctic foxes can adopt two different resource use strategies, referred to as coastal and inland. Interior habitats are more subject to seasonal variations in temperature influencing the availability of resources. The resident rock ptarmigan (*Lagopus muta*) and migrating birds like waders, geese and passerines make up for most of the inland foxes’ diet [16,19,25,32]. The Western part of Iceland bears the most productive seashores, with a greater productivity than northern, southern and eastern Iceland combined [33]. It also supports most of the large seabird colonies that nest on the cliffs during summer [25]. Ice-free shores contribute to the stability of food supply [34], enabling coastal foxes to benefit from carrion from marine mammals, fish and marine invertebrates in addition to seabirds and some terrestrial preys [19]. All foxes can occasionally consume sheep (*Ovis aries*) and reindeer (*Rangifer tarandus*) carcasses, as well as cattle and horse carcasses that are used as baits by hunters [16]. Because of their more stable food resources, coastal foxes are thought to be more territorial whereas inland foxes are more mobile [19]. This subdivision is reflected in genetic differentiation between coastal foxes from the north-western part of Iceland and the foxes from the rest of the country [35]. Because of this clear distinction, we carried out all analyses addressing coastal and inland foxes separately.

### Arctic fox samples

Fox mandibles were obtained from the collection of the Icelandic Institute of Natural History in Reykjavik, which consists of 12,200 mandibles from Arctic foxes culled from 1979 to present. Legally killed fox carcasses are donated voluntarily by hunters from all over Iceland. All foxes have been aged by counting annual cementum lines of canine tooth roots [36] at Matsons laboratory (United States). To include both coastal and inland foxes and assure a sufficient sample size, we chose jaws of foxes culled in Nordur Isafjardarsysla (henceforth NIS) and Nordur Múlasysla (NMU), two counties respectively representing the Western productive seashores and the Eastern inland areas (Fig 2). Individuals from NIS were culled within 5km from the coast and can thereby be qualified as coastal whereas foxes culled in NMU were located further than 50km from the coast and are thus defined as inland. We included 106 samples from [28] culled in the same counties, along with samples coming from 5 other counties whose ecology was either similar to NIS or NMU (Ester Unnsteinsdóttir, personal communication), and culled within 5km from the coast or further than 50km from respectively. Altogether, this added up to a total of 256 samples, 127 of them being from inland areas and 129 being from coastal areas, covering a period of 40 years, from 1979 to 2018. The last 14 years, characterized by large fluctuations in abundance, were sampled more intensively. All the foxes analyzed were between one and two years old. As previous research showed that resource use as inferred from δ^13^C and δ^15^N does in general not differ between sexes [32,37] male and female individuals have been chosen at random (S1 and S2 Tables).

**Fig 2.**
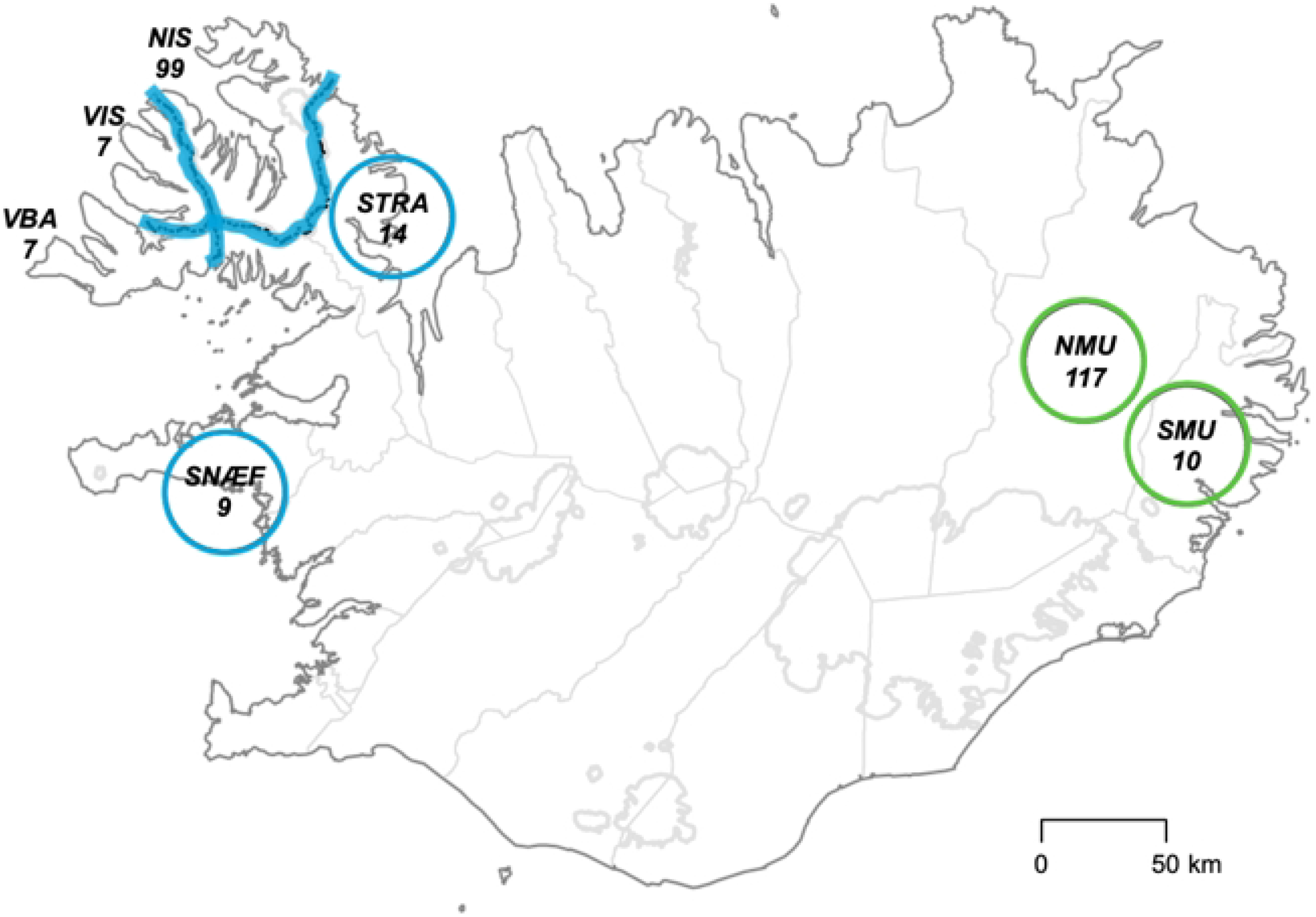
Map of Iceland displaying the different culling locations. Coastal areas are shown in blue, and inland areas in green. Culling locations are circled when hunting does not occur in the whole region. The number of foxes included in our dataset is specified under the abbreviation for each region.

The isotopic composition of one collagen has a slow turnover rate [38]. It can reflect a lifetime average dietary intake although it is biased towards the period of greatest growth [3], which is until 8-9 months old for Arctic foxes [25]. Here, we will assume that the isotope signatures are representative of the diet of Arctic foxes during their first year of life, thereby reflecting their average resource use during this period. Collagen has been extracted from lower jaws following the same protocol as [28], based on a standardized method from [39] and modified according to [40]. The samples were analyzed for stable isotopes of carbon and nitrogen at the Stable Isotopes in Nature Laboratory (SINLAB) at the Canadian Rivers Institute, University of New Brunswick.

### Prey samples

Greylag goose (*Anser anser*) muscle and egg samples along with northern fulmar (*Fulmarus glacialis*) muscle samples have been provided by the Icelandic Institute of National History. Both muscle and egg samples have been prepared for stable isotope analysis following the method from [41]. They have been analyzed for stable isotopes in carbon and nitrogen at SINLAB, along with fox collagen samples. Additional prey signatures of ptarmigan, common eider (*Somateria mollissima*), wood mouse (*Apodemus sylvaticus*), golden plover (*Pluvialis apricaria*), whimbrel (*Numenius phaeopus*), sheep, horse, kittiwake (*Rissa tridactyla*), starfish (*Asteria rubens*), redshank (*Tringa totanus*), common snipe (*Gallinago gallinago*), and black guillemot (*Cepphus grille*) were obtained from [28].

### Statistical analysis

The statistical analysis was performed using R version 4.0.4 [42]. Stable isotope ratios were expressed using the standard δ notation in parts per thousand (‰) [43]. The international standards, i.e the Vienna Peedee Belemnite for δ^13^C and atmospheric nitrogen for δ^15^N, were used as reference [43]. The distribution of Arctic foxes’ isotopic signatures with respect to their potential preys was assessed graphically by plotting individual values from both habitats. Prey values were corrected for isotopic discrimination, which corresponds to the amount of change in isotope ratios occurring as a prey is incorporated into the consumer’s tissue [43,44]. Since the discrimination factor for Arctic fox bone collagen has not been determined experimentally, we used the experimentally determined values from Arctic fox blood cells [37], modified according to [45], who estimated the fractionation between wolf red blood cells and bone collagen. Prey signatures have consequently been adjusted by 3.09 ± 0.25‰ and 3.36 ± 0.37‰ to account for diet to bone collagen discrimination of ^13^C and ^15^N, respectively (Fig 3). Both foxes and preys’ raw δ^13^C values have also been corrected for the Suess effec<t, which consists in a depletion in δ^13^C in the biosphere driven by the input of CO_2_ from fossil fuel since the Industrial Revolution [46]. We followed the same method as [28], i.e using a mean δ^13^C rate of change of −0.026% per year [47] and correcting all the δ^13^C isotopic ratios to levels, which correspond to the first year of the study (1979).

**Fig 3.**
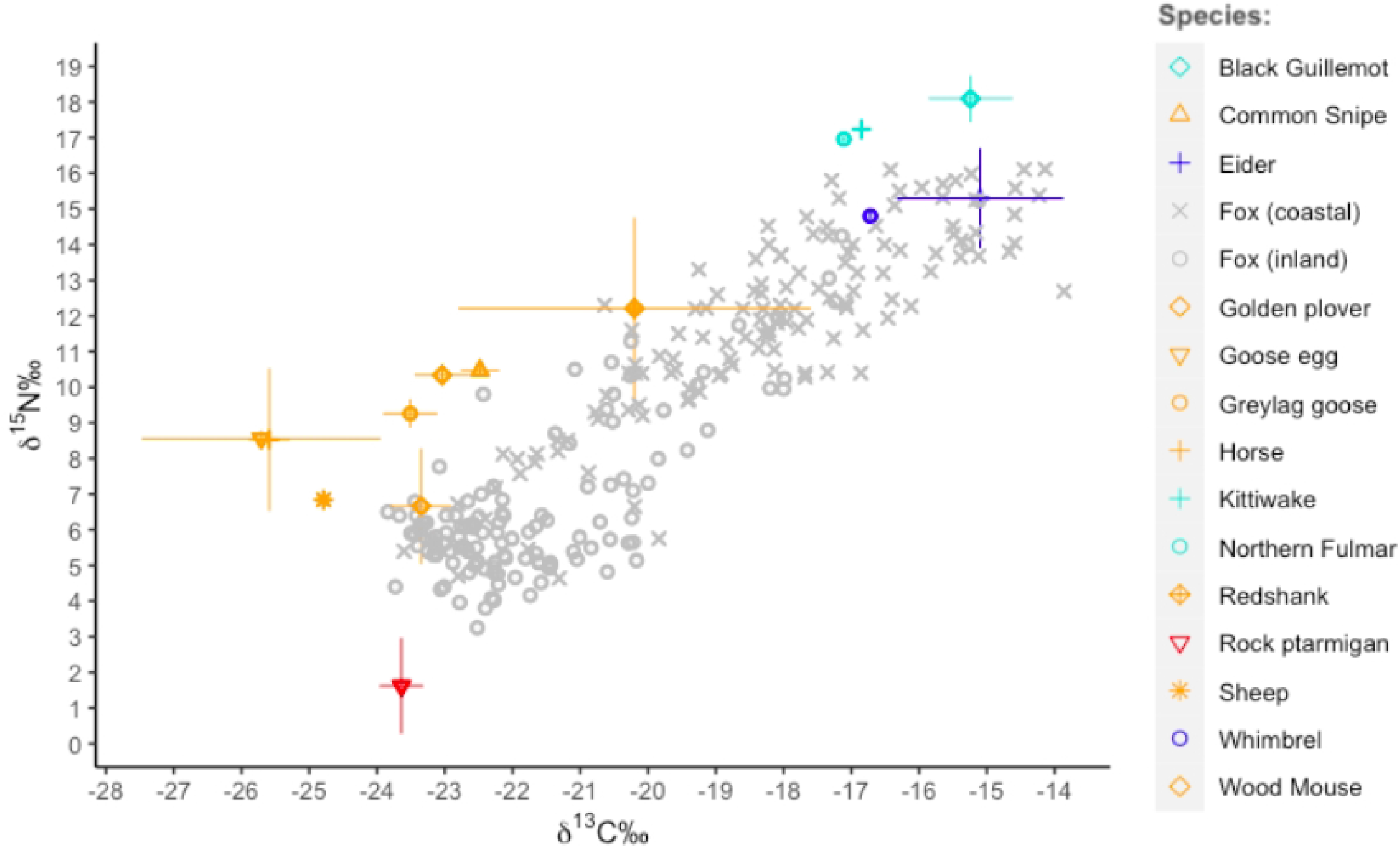
Isotopic signatures (‰) of foxes from coastal and inland habitats plotted along with their potential prey species corrected for trophic discrimination. Discrimination factor from Arctic fox blood [37], modified following [45], and plotted with their respective standard deviation.

Temporal changes in Arctic fox isotopic values were analyzed using generalized additive models (GAM) with the mgcv package to allow for non-linearity [48]. Changes in δ^13^C and δ^15^N have been modelled as a smooth function of the birth year of each individual (Fig 4). An interaction allowed to fit different changes over time for inland and coastal foxes, and their different means with respect to the two isotopes ratios were modelled as a fixed effect. We chose the restricted maximum likelihood method and used the default parameters of the package for both smoothing parameter and the number of basic functions. Both *gam.check* and *concurvity* functions were used to test for the fit of the model as recommended by [49] (S3 Table).

**Fig 4.**
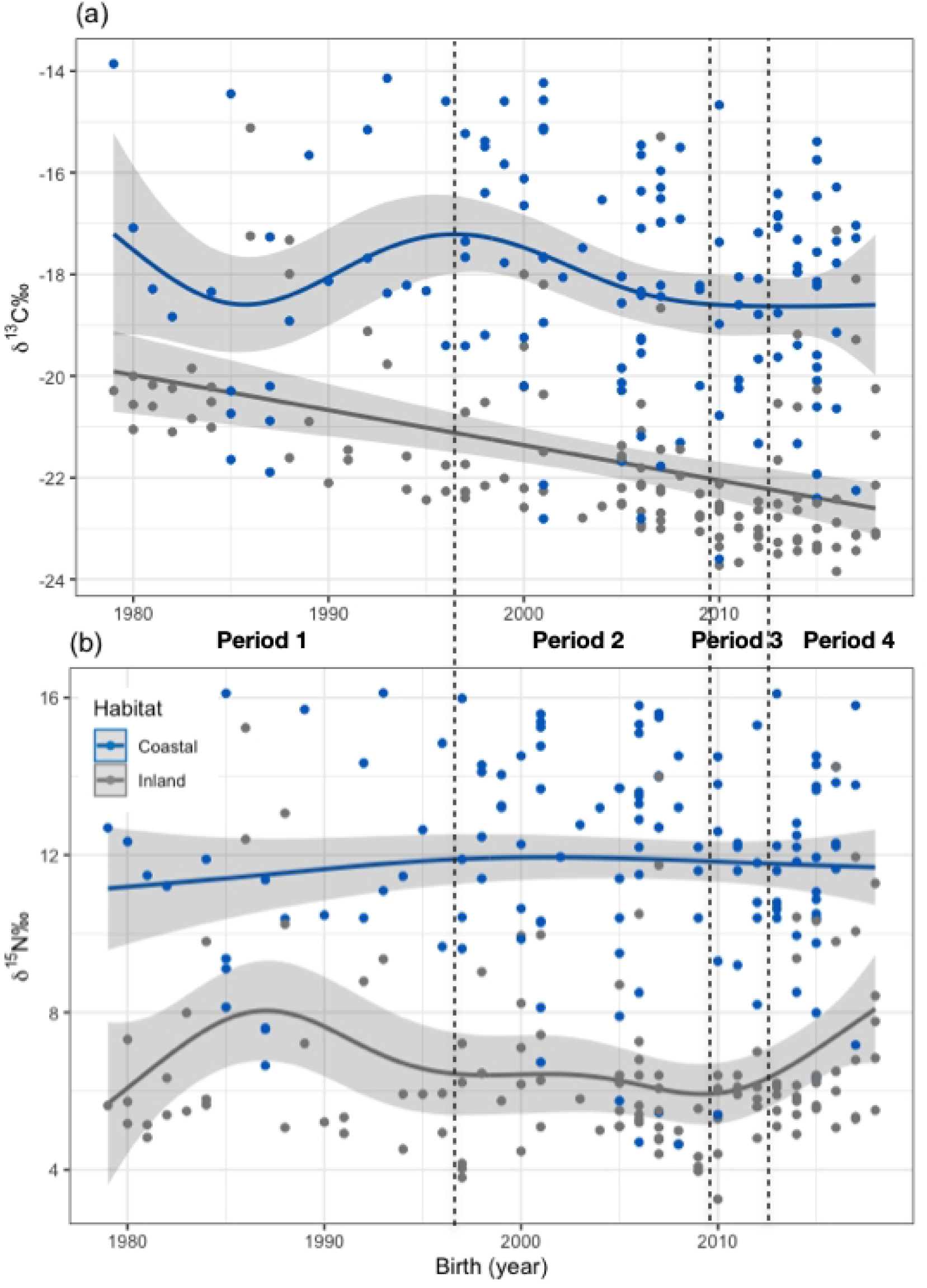
Isotopic signatures in (a) δ^13^C and (b) δ^15^N of coastal and inland foxes plotted according to their year of birth. Lines have been generated with generalized additive models, along with the 95% confidence interval.

The proportions of different prey in Arctic foxes’ diet over time were estimated using Bayesian stable isotope mixing models as implemented in the MixSIAR package [50]. As mixing models perform best with few potential sources, the different prey items were grouped considering the similarity of their isotopic signatures and their ecological relevance. We created four distinct groups: cliff nesting seabirds (black guillemot, northern fulmar and kittiwake) called *seabirds*, eider ducks that had distinct isotopic signatures from the other seabirds were grouped with whimbrels to *marine resources*, the rock ptarmigan was kept as a focal species, and all other terrestrial prey (common snipe, greylag goose, goose egg, redshank, golden plover, wood mouse, horse, sheep) were grouped to *terrestrial resources*. Because of their more marine signature, whimbrels were placed with eider ducks but the other wader species were placed in the terrestrial resources. To address potential changes in dietary composition, we defined four periods characterized either by growth or decline of the Arctic fox population (Fig 1). The mixing models were run separately for coastal and inland foxes. To account for the uncertainty considering the discrimination factor and because the results of mixing models can depend on how correct the discrimination value is [44], the analysis was repeated with the discrimination factor used in [28] (S4 Table). We ran the analysis following the MixSIAR manual recommendations, and did 300,000 MCMC replicates, (preceded by 200,000 burn-in) and used a residual*process error, as advised by [51]. The performance of mixing models being sensitive to the selection of priors [52], and as all priors are informative in a mixing context, we used priors based on the known dietary preferences of the foxes from both habitats, i.e. predominantly marine diet for coastal foxes and terrestrial diet for inland foxes [16,32]. Thus, the prior attributed to coastal foxes was 2/3 of marine diet components (1/3 seabirds and 1/3 marine resources) and 1/3 terrestrial diet components shared between the two groups (ie 1/6 rock ptarmigan and 1/6 terrestrial resources). For inland foxes the prior was opposite with 2/3 terrestrial diet components shared between the two groups and 1/3 marine diet components. The convergence of the MCMC estimations was assessed based on Gelman-Rubin and Geweke diagnostics, and we also inspected the correlation between different sources. In addition, we checked that the posterior distributions were unimodal. The isospace plots for the converging models have been added in S4 Fig.

For each period and habitat, we determined the isotopic niche breadth using the SIBER package [53]. The use of stable isotopes to infer a population’s trophic niche width is increasing as isotopic niches are considered to be a good proxy for ecological niches [54–56]. The ellipses being unbiased for sample size [53], it was possible to compare them even though the periods did not contain an equal amount of individuals.

## Results

The prey species showed the typical distinction between terrestrial species with lower δ^15^N and δ^13^C values and marine species with higher ratios for both isotopes (Fig 3). The fox values were within the polygon delimited by the prey and covered the whole gradient from terrestrial to marine resources. Many coastal foxes had signatures that placed them close to the marine prey, whereas inland foxes were in general placed at the other end of the coast-inland gradient. Accordingly, both isotopic ratios showed a significant difference between foxes of the two habitats, with an average difference of respectively 5‰ and 3‰ in δ^13^C and δ^15^N values between coastal and inland areas (Table 1).

**Table 1.** Parameter estimates from generalized additives models assessing the effect of birth year and habitat for (a) carbon isotopes (δ^13^C) and (b) nitrogen isotopes (δ^15^N) from bone collagen, along with their corresponding smooth terms values.

|  | Estimate | Std. Error | t value | P |
| --- | --- | --- | --- | --- |
| (a) $\delta^{13}\text{C}$ \| Formula: $\delta^{13}\text{C} \sim \text{s}(\text{Year}, \text{by} = \text{Habitat}) + \text{Habitat}$ | | | | |
| Intercept | 11.7720 | 0.2156 | 54.60 | <2e-16 |
| Habitat (inland) | -5.1445 | 0.3067 | -16.77 | **<2e-16** |
| (b) $\delta^{15}\text{N}$ \| Formula: $\delta^{15}\text{N} \sim \text{s}(\text{Year}, \text{by} = \text{Habitat}) + \text{Habitat}$ | | | | |
| Intercept | -18.7225 | 0.1629 | -111.88 | <2e-16 |
| Habitat (inland) | -3.4169 | 0.2301 | -14.85 | **<2e-16** |
|  | edf | Ref.df | F | p-value |
| (a) $\delta^{13}\text{C}$ \| Formula: $\delta^{13}\text{C} \sim \text{s}(\text{Year}, \text{by} = \text{Habitat}) + \text{Habitat}$ | | | | |
| Coastal | 4.338 | 5.329 | 2.472 | **0.0413** |
| Inland | 1.002 | 1.004 | 21.600 | **5.96e-06** |
| (b) $\delta^{15}\text{N}$ \| Formula: $\delta^{15}\text{N} \sim \text{s}(\text{Year}, \text{by} = \text{Habitat}) + \text{Habitat}$ | | | | |
| Coastal | 1.633 | 2.022 | 0.525 | 0.5835 |
| Inland | 4.542 | 5.562 | 1.889 | 0.0607 |

For each model, the fixed effect of the difference between habitats is given (Habitat (inland)) as well as the estimates for non-linearity (edf - effective degree of freedom) for foxes from each habitat.
Significant effects are shown in bold.

The GAM identified significant changes in δ^13^C over the study period. A clear linear decrease in δ^13^C values was observed for inland foxes (edf = 1.002; *p* < 0.001; Fig 4), while coastal foxes underwent non-linear variations (edf = 4.338; *p*= 0.041) with fluctuations in the start of the study period and more stable values in recent years. In contrast, δ^15^N ratios showed less temporal change. For coastal foxes, the values of δ^15^N were close to remain stable throughout the whole study period (edf = 1.633; *p*= 0.584; Fig 4) while more fluctuations were observed for inland foxes (edf = 4.542), with a noticeable increase at the end of the study period that was close to significant (*p*= 0.061). The *gam.check* function showed full convergence for both models, as well as randomly distributed residuals (*p*> 0.05 in both cases), thus confirming that the default parameters of the program were adequate. The *concurvity* function showed no evidence for concurvity between variables (S3 Table).

According to the results of the mixing models, the dietary composition of coastal foxes remained stable throughout the study period (Fig 5a) and the models converged well with respect to the discrimination factor chosen (S4 Table, S1 Fig). As expected, marine diet components dominated for coastal foxes, but contrary to our expectations seabirds were less important that other marine resources. However, the posterior diet proportions of seabirds and alternative marine resources were highly correlated (−0.94), indicating that the distinction between these two prey groups was difficult to estimate reliably. This was reflected in considerable overlap in the credibility intervals of the dietary proportions estimated (Fig 5a). Among terrestrial preys, rock ptarmigan were clearly the most important resource. Inland foxes had a more variable diet with respect to the different periods (Fig 5b) and the model converged with the discrimination factor (S4 Table, S1 Fig). Ptarmigans were always the most used resource, but their proportion varied over time, representing up to three fourth of the diet during the decline phase. Marine prey were more prominent during the two first periods of growth, while other terrestrial resources than the ptarmigan became increasingly important over time. The credibility intervals of seabirds did not exclude 0 in all periods, and this also applied to terrestrial resource during the two first periods. A sensitivity analysis was performed using the discrimination factor from [28], but no other model showed convergence for both coastal and inland habitats (S4 Table).

**Fig 5.**
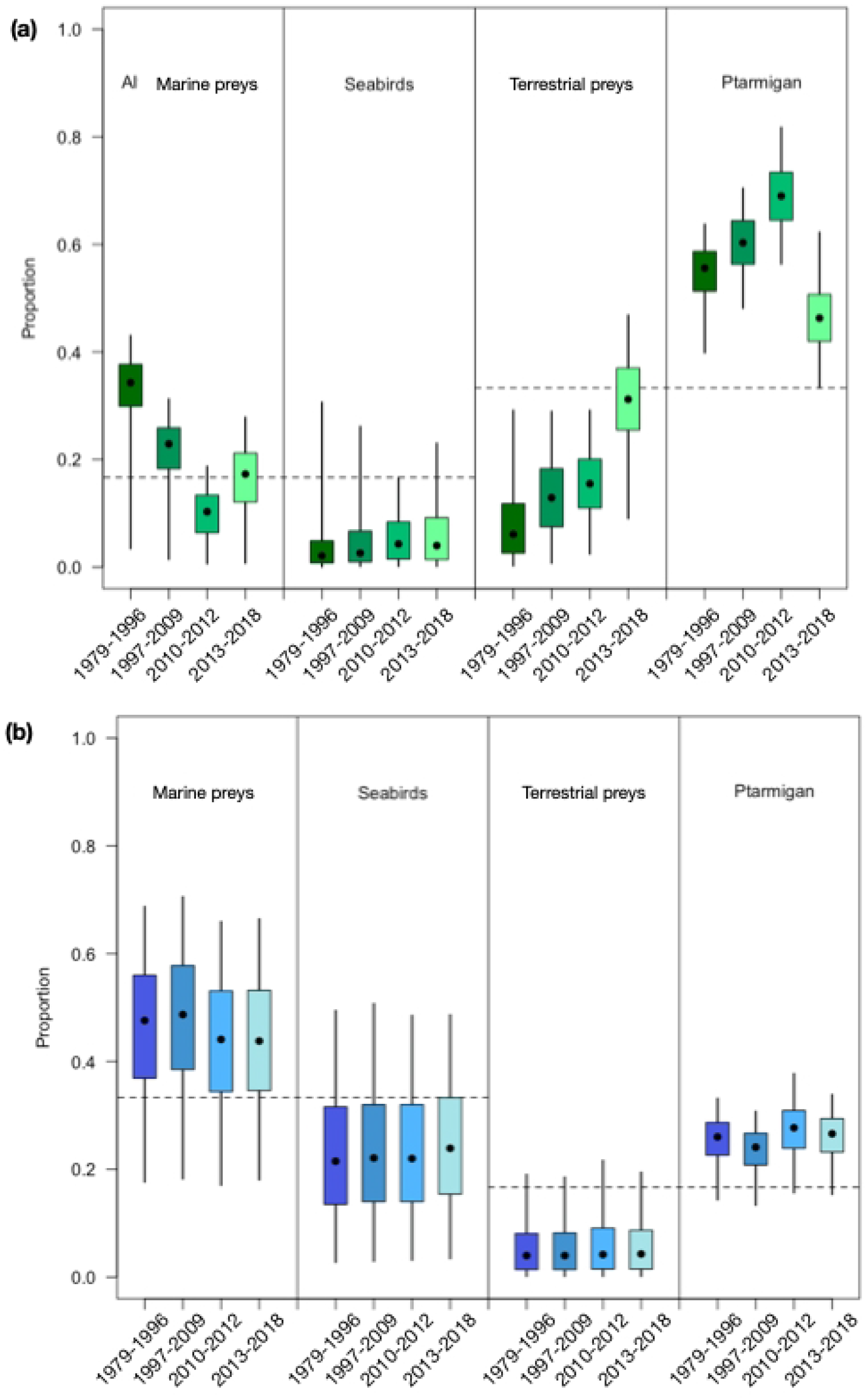
Box plots representing the proportion of different prey items in the diet of (a) inland and (b) coastal Arctic foxes during 4 different periods. The box plots are based on the results of the MixSIAR analysis. The median and the confidence interval are also represented.

As for the mixing models, the variations in the isotopic niche breadths were more important for inland foxes (Fig 6a). The niches shifted towards lower δ^13^C values over time and kept a similar width until the decline phase when the niche shrank considerably. Coastal foxes’ niches remained stable and overlapped during the different periods (Fig 6b). Moreover, the niche breadths of coastal foxes were overall greater than the ones of inland foxes (S2 Fig).

**Fig 6.**
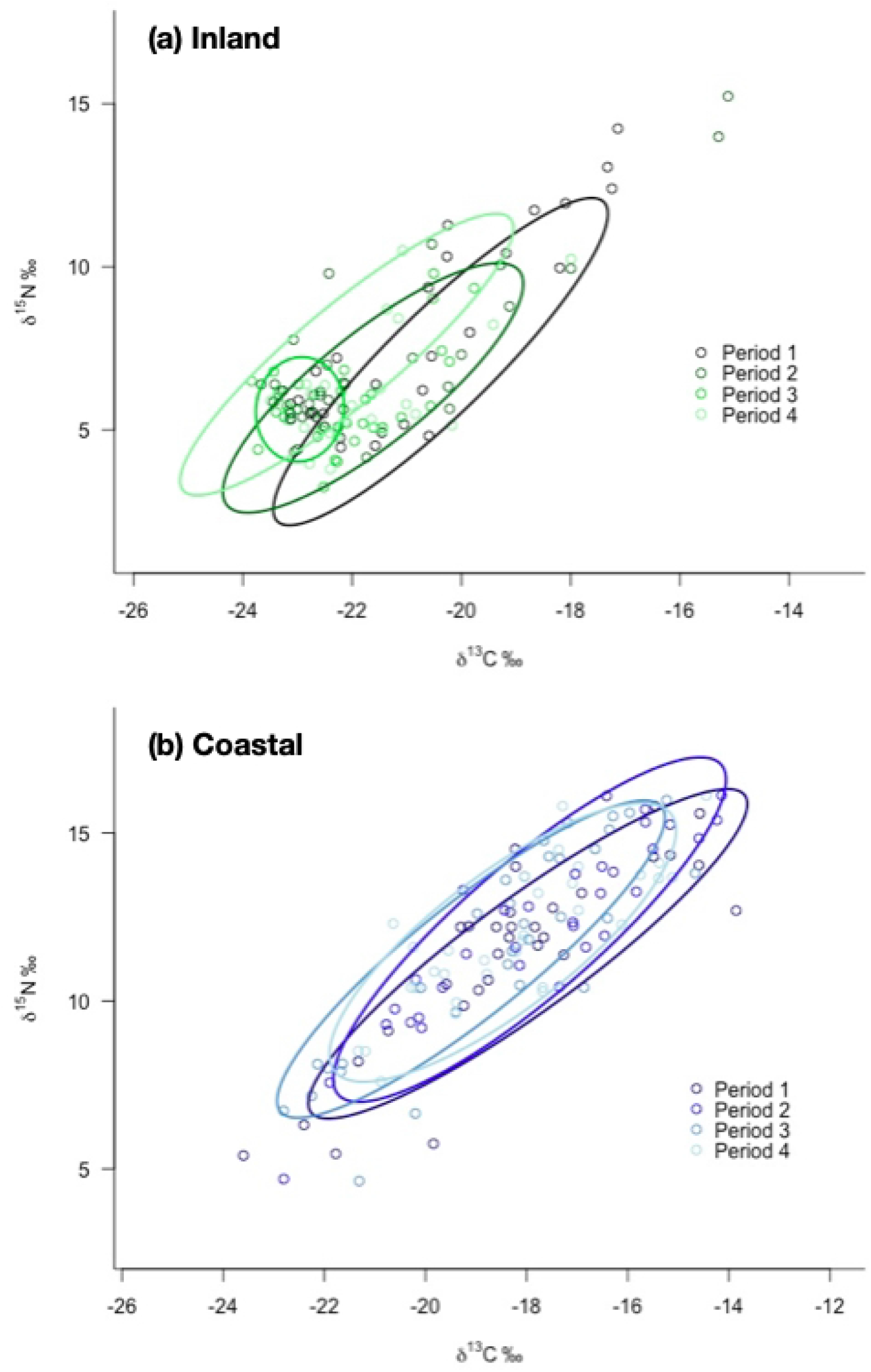
Standard ellipses representing the isotopic niches breadth from (a) inland and (b) coastal foxes during four different periods.

## 4. Discussion

### 4.1 Variations in resource use

Our results confirmed the previously described differences in diet between coastal and inland foxes, with coastal foxes having overall more marine isotopic signatures than inland foxes [28,34]. The significant decrease in the stable isotope ratio of carbon among inland foxes suggested a shift towards more terrestrial preys, and was combined with a slight increase in nitrogen ratios at the end of the study period. Coastal foxes, on the opposite, showed no statistical evidence for changes in nitrogen ratios, and the changes in carbon ratios they underwent during the beginning of the study had a low statistical support, and were likely due to the small number of individuals sampled during the first years of the study. These differences between a rather stable diet at the coast and more fluctuations in inland habitats were in agreement with the estimations of dietary composition from isotopic mixing models as well as the isotopic niche breadths, which showed important changes in the resource use of inland foxes while coastal foxes seemed to have a rather stable diet and constant niche width throughout the years.

In agreement with [28], who highlighted the importance of marine resources in the diet of Icelandic Arctic foxes, the mixing model analysis showed that marine preys were the main resource consumed by coastal foxes, and that they also were important to a certain degree for inland individuals. However, contrary to [28], we separated the marine resources in two groups and our results suggested that prey from lower trophic levels such as common eiders and whimbrels could be more important than cliff-nesting seabirds. Although the separation between the two groups of marine resources is not strongly supported by the mixing model (high correlation between the posterior distributions and overlap of credibility intervals), it appears likely that marine resources like common eiders, which are the most common waterfowl available throughout the year and are widely distributed in coastal areas, could represent an important part of the diet of Icelandic Arctic foxes. However, predation on eiders in Iceland is thought to be lower than for other populations in other areas since their protection is one of the reasons for the Icelandic fox culling program [26, 57]. A high importance of common eider in Arctic fox diet is also contrary to findings from prey remains at dens [20], where fulmars were the most common species. This unexpected result could possibly be due to the fact that many of the coastal foxes were culled at eider colonies (Unnsteinsdóttir, personal communication).

Alternatively, whimbrels, which were included in our marine preys, could have been an important resource for foxes from both habitats as they are abundant and accessible during the breeding season [58]. Unfortunately, the grouping used in the mixing models analysis makes it hard to determine whether or not this particular wader species was more important than the others, and no information was found in literature to either support or contest this assumption.

The rock ptarmigan, also a year-round resident in Iceland, appeared to be the preferred terrestrial resource. They were especially important for inland foxes and their increased proportion in the diet (Fig. 5b) may explain the observed decrease in carbon ratio over time (Fig. 4a).

During the breeding season, waders and geese are increasingly available to foxes as goose populations, in particular, are increasing [20]. In contrast with [20] who suggested that these resources were the main preys available to inland foxes, our results suggested that these prey items were of minor importance. This confirmed the results from [28] who did not find support for an increased use of geese, despite isolating the greylag goose as a focal source in their mixing model analysis. However, the latest period of growth in fox population size, period 4, showed a clear increase in the use of other terrestrial resources than ptarmigan among inland foxes. A shift from ptarmigan to other terrestrial resources such as geese, waders and their eggs is in agreement with the observed increase in δ^15^N observed for inland foxes in the end of the study period (Fig. 3b). In addition, the lack of species such as the pink-footed goose in our prey signatures could explain the apparent minor importance of this group since this particular species has been shown to be especially important for Iceland Arctic foxes [16,20].

### 4.2 Driver of population change

As suggested by [22], the fluctuations in the population size of Arctic foxes likely result from changes in carrying capacity due to changes in abundance of main resources. Accordingly, the results of [28] showing a constant and important use of marine resources indicated that increasing populations of seabirds could have been a major driver of the long population increase of the foxes from the 1980’s. For coastal foxes, our results showed a stable diet composition over the whole study period, despite considerable fluctuations in population size (Fig. 1). This is also consistent with a bottom-up regulation of the fox population. Indeed, after a period of increase for many seabirds in Iceland up to the turn of the millennium, populations stabilized and some dramatically declined during the last decades [59,60] when the numbers of foxes hunted in coastal areas started to fluctuate considerably. Breeding failure of several species was observed in 2005, while in 2010-2011 Puffin (*Fratercula arctica*) reproduction failed totally [61]. This was attributed to a lack of sandeel (*Ammodytes marinus*), a major resource for many seabird species [59,62] notably for Fulmars [63], a key prey of Arctic foxes [20].

The strong decrease in fox numbers in 2011-2012 that was more pronounced for the coastal population might be a direct consequence of this event. Moreover, our results suggested that Common Eider is likely to be an important prey for Arctic foxes. Their populations stabilized and partly declined in Iceland in the 2000s after an overall increase in the end of the last century [64] and notably declined in Westfjords in western Iceland after 2000 [65]. This resource may thus also have contributed to flattening out and periodic decline of the coastal fox population.

For inland foxes, Ptarmigan were the main resources during the whole study period, but their population trends cannot really explain the increase in the Arctic fox population as many Icelandic populations showed declining trends in the last decades [66]. As suggested by [28] it is likely that the increase in marine resources also resulted to an improved situation for inland foxes during the growth period. This is supported by the fact that the crash of seabird populations together with declining Eider populations in 2010-2011 led to an abrupt decline in foxes hunted in the inland as well, although the decline was not as strong as for the coastal population. Notably fulmars were indeed breeding on cliffs far inland and preyed upon by Arctic foxes, but at present, these inland colonies are less active (ER Unnsteinsdóttir, personal observations). During these years, the marine input in the diet of the foxes declined abruptly and ptarmigan dominated. However, according to [66] ptarmigan populations were in the low phase in these years, and could thus not compensate the lack of marine resource for the foxes. After this decline phase, inland foxes switched to use other terrestrial prey. This is in accordance with an increasing importance of geese and waders that experience positive population trends in inland areas [20]. In recent years the inland foxes thus adapted their diet to the changing resource situation shifting from declining Ptarmigan and seabirds to other terrestrial prey exhibiting a generalist strategy.

### 4.3 Population isotopic niche breadth

The high hunting pressure in Iceland leads to a high turnover in territorial foxes, and [22] suggested that Icelandic Arctic foxes engage in contest competition as they adapt their territory size in response to variations in carrying capacity.

Consequently, we predicted that Arctic foxes’ niche breadth would vary over time, but found no support for this hypothesis in the coastal habitat. Although the diet of coastal foxes seemed not to vary over the study period, one could have expected that the decline in the availability of seabirds would have led to a narrower niche breadth or to a shift in the diet. The apparent consistency in their isotopic niche at the population level could hide some variations at a finer scale - the individual scale.

The results from isotopic niche analysis suggested that coastal foxes globally have broader niches than inland foxes. In previous research, [34] pointed out the same phenomenon and suggested that these wider niches were due to a diversification of individual strategies likely dictated by the local abundance of resources. The heterogeneity of coastal areas could lead to increased individual specialization of some foxes, especially since coastal foxes are more territorial [19]. This assumption would support that the fluctuations in carbon ratios observed among coastal foxes are likely to be influenced by individual variations in the diet rather than by a global shift in resource used at the population scale. This specialization would be a way of reducing the potential dietary overlap among foxes, in response to an increasing intra-specific competition pressure [67].

Inland foxes on the contrary showed more variations in isotopic niche space over time, as well as a marked reduction in their niche breadth during the period of decline. This fits with the increased use of rock ptarmigan observed in the mixing model results during the decline period, but also with the assumption that this specific prey item, potentially in combination with other preys with low carbon signature like goose eggs, might have partly driven the decline in carbon ratio observed among inland foxes. In previous research, [19] even showed that the number of occupied fox dens was positively correlated to ptarmigan abundance, highlighting the importance of this prey item.

The reduction in the niche space matches the years of low productivity of seabirds, and illustrates the low availability of these resources, thus narrowing the niche. As the fox population recovered, the niche size widened, which could indicate that some marine species were available again, and were combined to an increasing use of other terrestrial resources.

## 5. Conclusion

Both marine and terrestrial ecosystems in Iceland are at present changing under the direct and indirect effects of climate change. Our results showed how the Arctic fox, a generalist top predator that uses different resources in western (coastal) and eastern (inland) Iceland, reacted to changes in resources availability. Coastal foxes that benefit from the productive seashores of western Iceland exhibited a constant marine dominated diet over the 40 years of our study. When seabird populations experienced reproductive failure, the fox population declined, probably because there were no alternative resources accessible. Arctic foxes are indeed genuine generalists able to exploit a wide variety of resources. As the seabird species, the foxes are experiencing the profound changes in the marine food web related to lower reproduction of the sandeels that is related both to ocean warming and to structural changes of the marine food web [59,61]. Inland foxes on the contrary changed their diet and likely increased their use of terrestrial prey whose populations are increasing, such as geese and waders. From being a stronghold of the Icelandic Arctic fox, with climate change the coastal areas may become a habitat with less reliable resources, whereas the prey basis in the inland areas is becoming more productive.

## 5. Acknowledgements

The authors would like to thank Sissel Kaino from the Biology laboratory of the Arctic University of Norway for her help in the lab.

## 7. Supporting information

**S1 Table. Year of birth, location and sex of the foxes used in this study.**

**S2 Table. Extractions**. Extraction of [28] along with the one carried out in the present study.

**S3 Table. Outputs from (a) *gam.check* and (b) *concurvity* functions to test for the fit of the generalized additive models**. Estimated with the default parameters of the *gam* function in the mgcv package.

**S4 Table. Overview of the runs performed in MixSIAR.** Foxes were analyzed separately depending on their habitat and time was included as a categorical (four periods) covariate. Coastal and inland foxes were analyzed separately. We used two different discrimination factors. The first one based [28], and the second one based on the values from Arctic fox blood [37], modified according [45]. Convergence was assessed based on the Gelman-Rubin (Gel.) and Geweke (Gew.) diagnostics. The runs that did not converge or that did not result in unimodal posterior distributions for dietary proportions (Unim.) were discarded. The maximal correlation between two sources is given (Cor.). Sources are abbreviated as Ptarmigan – P, Alt. terrestrial – T, Alt. Marine – M, Seabirds – S. Mean posterior estimates of dietary proportions are given with 95% credibility intervals for the models which had satisfactory convergence diagnostics (in bold).

**S1 Fig. Isospace plots generated by MixSIAR on the convergent runs for (a) coastal and (b) inland habitats.** The discrimination factor used for this model was based on the values from Arctic fox blood [37], modified according to [45].

**S2 Fig. Standard ellipse area determined with the SIBER package for (a) coastal and (b) inland foxes over the four periods.**

